# *CompBioAgent*: An LLM-powered agent for single-cell RNA-seq data exploration

**DOI:** 10.1101/2025.03.17.643771

**Authors:** Haotian Zhang, Yu H. Sun, Wenxing Hu, Xu Cui, Zhengyu Ouyang, Derrick Cheng, Xinmin Zhang, Baohong Zhang

## Abstract

Advancements in high-throughput biological technologies, particularly single-cell RNA sequencing (scRNA-seq), have generated vast amounts of complex data that offer valuable insights into gene expression, cellular behavior, and disease mechanisms. However, the analysis of such data often requires specialized computational skills and expertise, which can pose a barrier to many researchers. To address this challenge, we developed CompBioAgent, a user-friendly web application designed to democratize access to bioinformatics resources, powered by Large Language Models (LLMs). By integrating with the CellDepot, CompBioAgent allows users to easily query and explore gene expression data related to various diseases, cell types, and experimental conditions. Moreover, the tool employs Cellxgene Visualization In Plugin (Cellxgene VIP) platform to generate a range of intuitive visualizations, including violin plots, UMAP embeddings, heatmaps, etc. With its natural language interface, CompBioAgent automatically converts scientific queries into structured data requests and returns the plot of interest, enabling seamless exploration of biological data without the need for programming expertise. Such integration enhances the ability of researchers to visualize complex biological insights and identify key patterns, making high-level data analysis more accessible and efficient. A demo website and a list of examples are available at https://celldepot.bxgenomics.com/compbioagent, and the source code is released at https://github.com/interactivereport/compbioagent.

## 1 Introduction

Single-cell RNA sequencing[1] (scRNA-seq) has revolutionized transcriptomic profiling by enabling the analysis of gene expression at the individual cell level. The extensive scRNA-seq datasets[2, 3] can be queried in various databases, such as CellDepot[4], which houses over 270 datasets from eight species and various tissues. In addition to CellDepot, several other public databases have been developed to facilitate the exploration of scRNA-seq data with ease. To name a few, the Single Cell Portal[5], hosted by the Broad Institute, provides access to a wide array of scRNA-seq datasets, including those from the Human Cell Atlas[6] and other consortia. It offers built-in exploration functions, such as t-SNE or UMAP embeddings and violin plots for visualizing gene expression across cell types and clusters. Similarly, the Single Cell Expression Atlas[7] hosted by EMBL-EBI serves as a public repository of single-cell gene expression data, allowing users to search across multiple species and studies.

To facilitate the analysis of these complex datasets, tools like Scanpy[8], Rapids-singlecell[9], and ScaleSC[10] offer functionalities for preprocessing count data, dimensional reduction, identifying marker genes, and visualizing data through methods such as t-SNE and UMAP. These tools have been instrumental in advancing complex data interpretation by providing comprehensive workflows for single-cell RNA-seq data analyses.

To further enhance the exploration and visualization of scRNA-seq data, web-based tools like Cellxgene[11] and Cellxgene VIP[12] have been developed for much easier exploration through user-friendly web interfaces. Cellxgene is an interactive data explorer for single-cell datasets. Such as those from the Human Cell Atlas, it integrates multiple tools to enable fast visualizations of at least 1 million cells, helping biologists and computational researchers explore their data effectively. Building upon this, Cellxgene VIP extends the capability of the original Cellxgene platform, offering over eighteen commonly used quality control and analytical plots in high resolution with highly customizable settings. Additionally, it provides advanced analytical methods such as marker gene identification, differential gene expression analysis, and gene set enrichment analysis. Cellxgene VIP also pioneers methods to visualize multi-modal data, including spatial transcriptomics and multi-omic datasets.

Despite the availability of these tools, many scientists face challenges in effectively utilizing them. To address this gap, the development of chatbots that integrate the power of large language models (LLMs)[13, 14] with scRNA-seq data analysis is essential. LLMs, such as GPT[15], and Llama[16] have demonstrated remarkable capabilities in understanding and generating human-like text, making them valuable assets in various domains, including bioinformatics. By leveraging LLMs, it is possible to create intuitive applications that allow users to query and analyze scRNA-seq data using natural language[17, 18, 19, 20, 21], thereby democratizing access to advanced analytical tools. The integration of LLM-powered agents with platforms like CellDepot can significantly enhance the accessibility and usability of scRNA-seq data analysis. In this study, we proposed CompBioAgent, a new tool that can interpret user inputs, generate visualizations, and provide insights without the need for any programming knowledge. This approach not only streamlines the analytical process, but also empowers a broader range of scientists to engage with complex scRNA-seq datasets effectively.

## 2 Methods

### 2.1 Overview of CompBioAgent

The primary idea of CompBioAgent is to enable users to easily interact with scRNA-seq data by natural language queries for exploration and visualization. CompBioAgent efficiently converts user’s queries into structured commands in JSON format, which can be further used to query relevant datasets and generate visualizations based on it.

### 2.2 Prompt engineering and JSON formatting

The first step in the CompBioAgent pipeline involves converting natural language into structured JSON commands. Using prompt engineering techniques, CompBioAgent parses the user’s input by leveraging large language models (LLMs) and well designed prompts to identify key elements such as gene names, diseases, cell types, experimental conditions, and desired visualization outputs. Through two types of prompt templates: complex and simple, CompBioAgent is able to transform natural language into well-structured JSON outputs through adding prompts right before the initial user inputs. The complex version provides detailed instructions for extracting information from queries, which leads to a more accurate conversion. More details can be found in the supplementary material. While the simple version focuses on more straightforward queries as shown below, optimized for higher efficiency and low cost.

**Prompt (simple): general instruction**

To convert a customer’s query into a JSON output for CellDepot API, please use the following guideline:

1. **Question**: Include the customer’s query.

2. **App**: Always set to “CellDepot”.

3. **Query**: Includes:

- **Database**: Always set to “project”.

- **Disease**: Match the disease to known names or list as entered.

- **Experiment type**: Default is [“scRNA”, “scRNA”].

4. **Action for query results**: Specify how the results should be displayed:

- **scRNA-Seq**:

- **App**: Always set to “CellDepot”.

- **Plot**: Select from embedding plot, violin plot, dot plot, stacked barplot, or heatmap based on the query details.

- **Plot options**: Include relevant options based on the plot type.

\### JSON Fields by Plot Type

1. **Violin Plot** (default for one gene):

- **Gene**: The gene mentioned.

- **Group by**: Default is “Cell Type”.

- **Sub-group by**: Use if the query specifies, e.g., “Disease”.

2. **Embedding Plot**:

- **Gene**: The gene mentioned.

- **Annotation**: Default is “Cell Type”.

- **Split by**: Default is “Disease”.

- **Embedding layout**: Default is “umap”.

3. **Stacked Barplot**:

- **Annotation**: Default is [“Cell Type”, “Disease”].

- **Color by**: Default is “Cell Type”.

- **Layout**: Default is “proportion”.

4. **Dot Plot** (default for two or more genes):

- **Cell selection**: Use to include/exclude cells based on metadata.

- **Gene**: List of genes mentioned.

- **Group by**: Default is [“Cell Type”, “Disease”].

5. **Heatmap**:

- **Annotation**: Default is [“Cell Type”, “Disease”].

- **Gene**: List of genes mentioned.

Both versions maintain the same standardization rules, including:

- Disease name normalization (e.g., “RRMS” or “PPMS” are converted to “Multiple Sclerosis (MS)”)
- Gene symbol standardization (e.g., “Tumor Supressor P53” to “TP53”, “CREB-1” to “CREB1”)
- Cell type standardization (e.g., “Micro” to “microglia”, “Astro” to “astrocytes”)
- Automatic plot type selection based on query content

Even with a specific prompt, a language model (LLM) may struggle to generate the correct JSON output. To address this, Few-shot prompting[22, 23] is introduced, a technique where a small number of examples are provided to guide the model’s response to a particular task. This method lies between zero-shot learning (with no examples) and fully supervised fine-tuning (which requires large labeled datasets). By including a few examples along with their corresponding chain-of-thoughts in the prompt, CompbioAgent can effectively process various user inputs and return the correct JSON output.

**Few-shot prompting**

Here are the instructions on how you are going to help customer use the web application, for any input from the customer, e.g.

Customer: Give me the gene expression of TYK2 in MS disease separated by disease vs. control You would be forming a JSON file like:

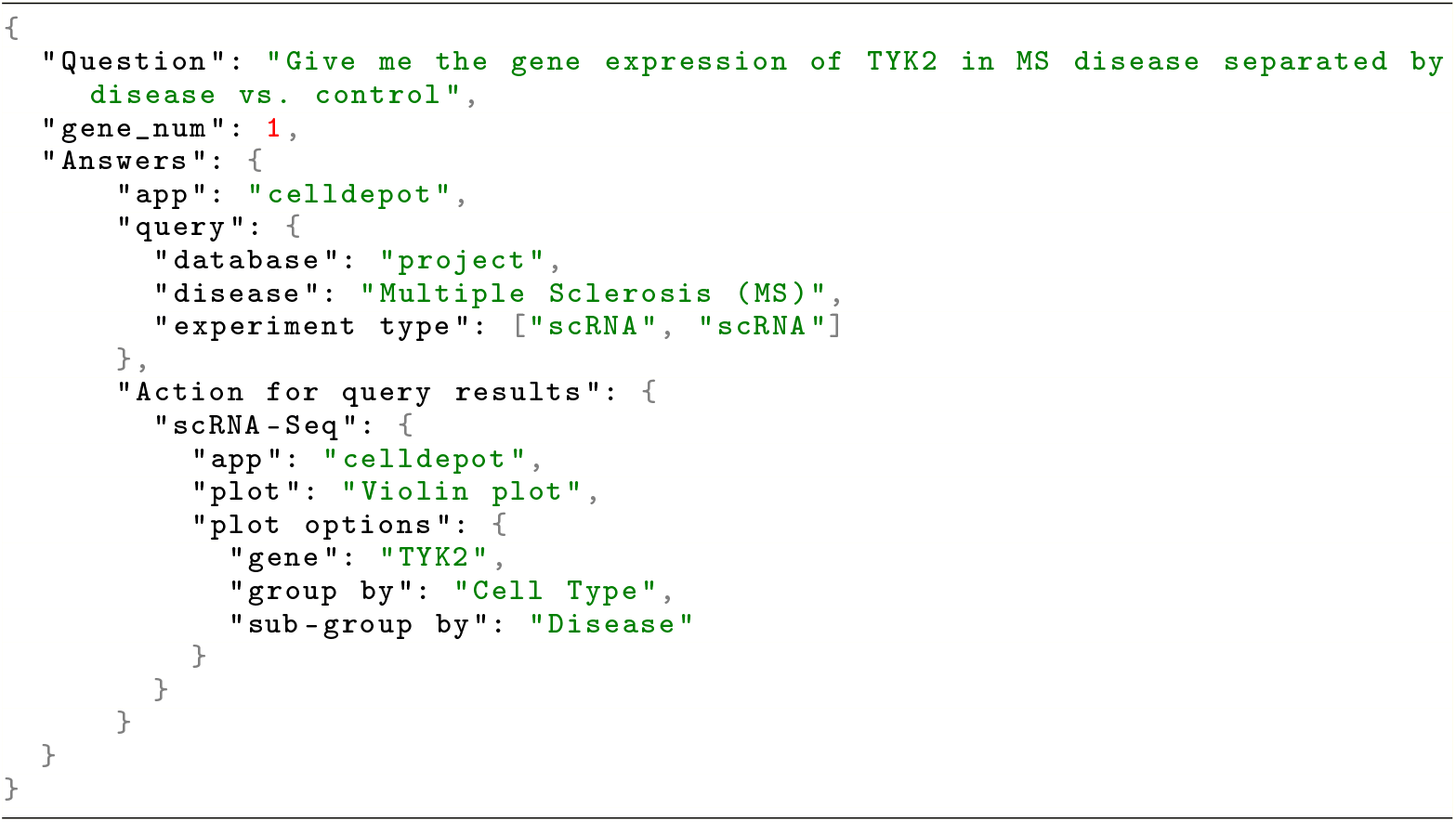

The idea above is to find a project in CellDepot that is related to the disease Multiple Sclerosis (MS), and display the expression of gene TYK2 between disease vs control in each cell type.

Here are more details about the JSON file fields.

-Question, repeat the questions asked by the customer.

-gene_num, the number of genes asked by the customer.

-app, the application that will use the JSON file. At this time the app is always CellDepot. In the future we will add more applications.

-query, the search condition sent to the application.

-(within query) database: allowed choices are application-specific. In CellDepot, the choices are project or gene. In our initial examples, the database field will always be project. In the future we may use gene for the database field.

-(within query) disease: In this example, we match MS from customer question to Multiple Sclerosis (MS).

-(within query) experiment type: the allowed values for experiment type for CellDepot are: scRNA, scRNA, Spatial Transcriptomics, CosMx. The default is to choose both scRNA and scRNA.

-Action for query results, how the application should show the data based on query results.

-(under Action for query results) scRNA-Seq: the type of action. The allowed values include scRNA-Seq, Bulk RNA-Seq, etc. For CellDepot, it is always scRNA-Seq at this point.

-(under scRNA-Seq) app, at this time the app choice for scRNA-Seq is always CellDepot.

-(under scRNA-Seq) plot, the possible plot values for scRNA-Seq are as listed before: embedding plot, violin plot, dot plot, Stacked Barplot, heatmap. In this case customer gives one gene in the question, so we use violin plot.

-(under scRNA-Seq) plot options, the options for the plot type selected above. In this example, violin plot.

-(under plot options) gene, this is a required field for violin plot, the gene asked by the customer in the question.

-(under plot options) group by, how cells are grouped, use one metadata field; the default choice for violin plot is cell type. Other possible choices are disease, treatment, sample, etc.

-(under plot options) sub-group by, how cells in each group are further divided, use one metadata field. Since the customer is interested in disease vs control, use disease field for this option.

For example, a query such as “Show me the expression pattern of neurotrophic factor genes in neurons and astrocytes in Alzheimer’s Disease” is parsed and transformed into a JSON structure containing essential information, including the disease name (Alzheimer’s Disease), the cell types (neurons, astrocytes), the genes of interest (BDNF, NGF, NTF3, NTF4), and the desired type of visualization plots (dot plot). More detailed formulations are shown in Results.

The version of prompt selection can be specified in the JSON through an optional ‘version’ parameter. If not specified, CompBioAgent defaults to the ‘complex’ version working with more advanced models like GPT-3.5, GPT-4o. If the ‘simple’ version is chosen, locally deployed smaller models like Llama 3 will be used.

CompBioAgent now supports five types of visualizations: embedding plots (UMAP/t-SNE), violin plots, dot plots, stacked barplots, and heatmaps. The prompt incorporates logic to automatically select appropriate visualization types based on the query content, and more details are shown in Section 2.3. As an example, the query “Compare gene SOD1 expression between MS disease patients and normal subjects” automatically selects a violin plot, while the query “Show the expression of SOD1 and TARDBP in MS” defaults to a dot plot.

### 2.3 Database querying and information extraction

Once the input is converted into the structured JSON format, CompBioAgent queries the appropriate databases, such as CellDepot, for the extraction of relevant datasets. The query information in the JSON file helps CompBioAgent precisely locate to the correct database, disease category, cell type and experimental condition. Currently, we demonstrate our application using several public datasets that are available in CellDepot, covering the following diseases:

- Multiple Sclerosis (MS)[24]
- Glaucoma[25]
- Diabetes[26]
- Systemic Lupus Erythematosus (SLE)[27]
- Alzheimer’s Disease (AD)[28]
- Cutaneous lupus erythematosus (CLE)[29]

CompBioAgent also supports a wide array of common cell types typically found in neurological and immune systems. These include neurons, astrocytes, microglia, and oligodendrocytes, as well as oligodendrocyte precursor cells (OPC) and ependymal cells. Immune cell types such as B cells, T cells, macrophages, and natural killer cells are also included, along with phagocytes and stromal cells. This comprehensive range of cell types allows for detailed exploration of cellular dynamics across various tissues and conditions.

### 2.4 Plot generation and visualization

Once the relevant data is extracted, CompBioAgent generates visualizations based on the user’s query. Supported visualizations include UMAP/t-SNE embeddings, violin plots, dot plots, stacked bar plots, and heatmaps. For instance, when a user queries a single gene of interest, CompBioAgent defaults to generating a violin plot, whereas multi-gene queries are visualized as dot plots. The automatic selection of visualization types ensures that the generated plots are tailored to the specific queried data and the nature of the user’s query.

Once the plot type is determined, CompBioAgent automatically invokes CellxgeneVIP, a powerful visualization tool, to generate the requested plots based on the queried dataset. CellxgeneVIP provides advanced interactive capabilities and an user-friendly interface for creating high-quality visualizations. The internally seamless connection to CellxgeneVIP enhances the plotting process by leveraging the tool’s sophisticated analytical functions. This includes a range of visualization options, from gene expression patterns in different cell types to the exploration of complex gene interactions. CompBioAgent’s ability to parse user queries, select the appropriate dataset, and invoke CellxgeneVIP for real-time visualizations ensures that users receive highly informative, accurate, and visually appealing outputs.

### 2.5 Integration with Large Language Models (LLMs)

To enable model accurately understanding user’s questions, CompBioAgent integrates both commercial and open-source large language models (LLMs), including OpenAI’s GPT models, Anthropic Claude models, Groq models, DeepSeek R1 models via API, as well as the local Llama 3 model powered by Ollama. These models are used to interpret natural language queries and generate the corresponding JSON output for querying the database and creating visualizations.

### 2.6 Deployment and error inspection

CompBioAgent is distributed as a web server powered by Apache and MySQL, supporting both CPU-only and GPU-enabled deployments. Typically, GPU is not necessary when user choose commercial LLM API. However, when user prefer not to expose their data to public APIs, Ollama is required and need to be properly configured in local and a NVIDIA GPU is suggested for speeding up token generalization.

During test, We have observed that ChatGPT engines do not always return consistent responses, with an approximate 5% chance that the LLM engine may provide incorrect outputs. For instance, when users request a list of the top 10 genes associated with a particular disease, the LLM engine might return placeholder terms such as TOP10_GENE_1, TOP10_GENE_2, instead of actual gene symbols like CREB1, WAS7P, etc. Attempting to plot such data would result in an error from the plotting tool. Similarly, in cases where UMAP plots require two columns—disease and cell type—the LLM engine may sometimes return only one column, leading to a failure in the plot generation.

To address these issues, we have implemented robust error handling and self-correction mechanisms. If inconsistent or incomplete responses are detected, an error message will prompt the user to try a different LLM engine. Furthermore, we have introduced additional checks to automatically fill in missing information when possible, reducing the likelihood of errors and improving the overall user experience. These measures ensure that CompBioAgent can effectively handle potential failure modes, such as inconsistent responses or missing data, without compromising the functionality of the tool.

## 3 Results

### 3.1 JSON formatting

Here we present two examples demonstrating how CompBioAgent formats JSON commands based on user’s input for querying and visualizing single-cell RNA-seq data:

#### Example 1: Single-Gene Expression in Alzheimer’s Disease

Query: “Show the expression of neurotrophic factor genes in neurons and astrocytes in Alzheimer’s Disease.”

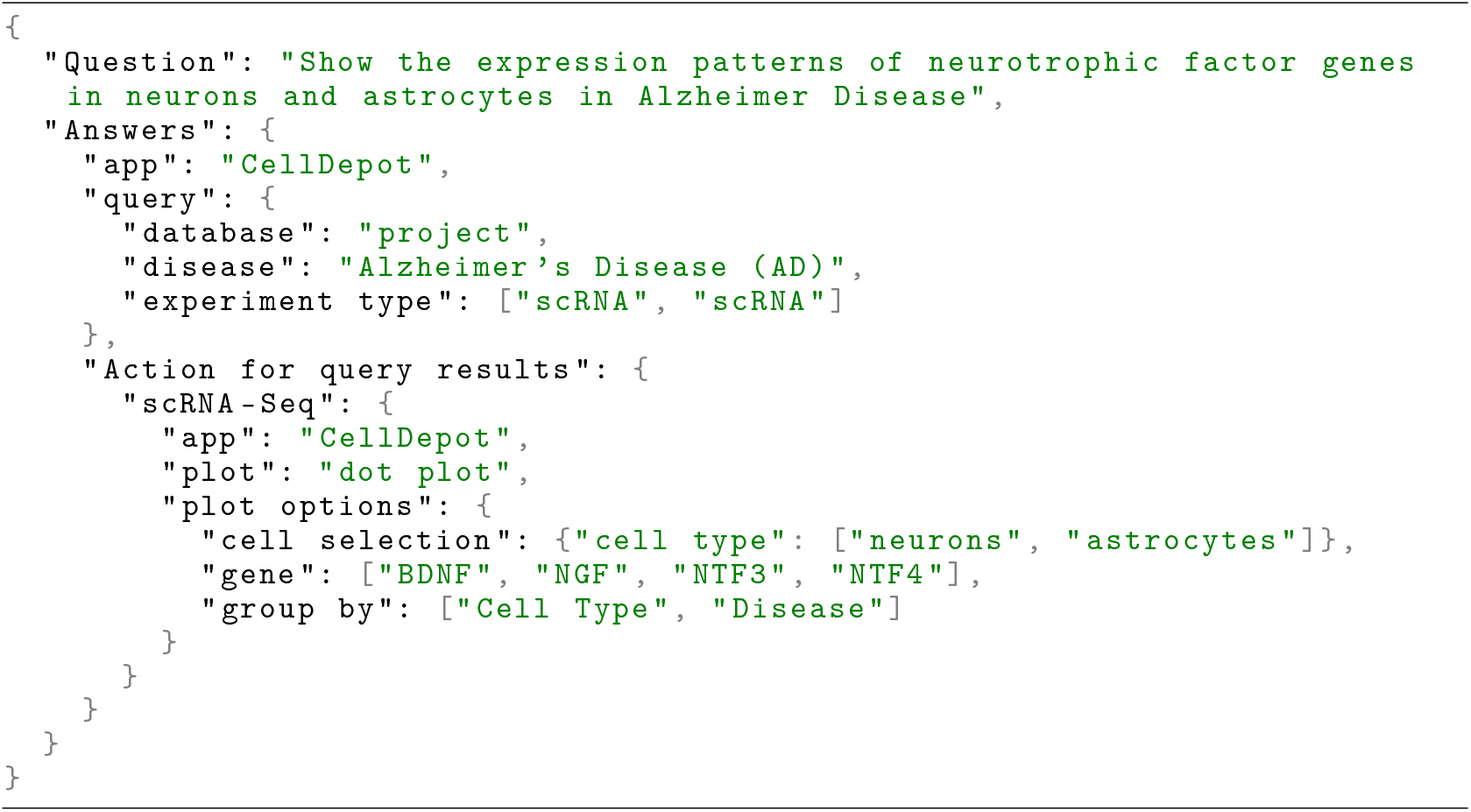

The goal of this example is to query CellDepot for a project related to Alzheimer’s Disease (AD) and visualize the expression patterns of neurotrophic factor genes (such as BDNF, NGF, NTF3, and NTF4) in specific cell types: neurons and astrocytes. The visualization is generated as a dot plot, with data grouped by both cell type and disease condition.

#### Example 2: Single-Gene UMAP plots with cell types in MS Disease

Query: Show TYK2 UMAP plots in MS, separate the plot by disease conditions, and show cell types.

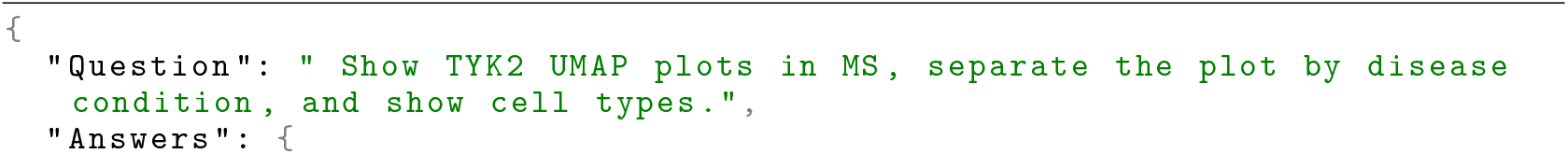
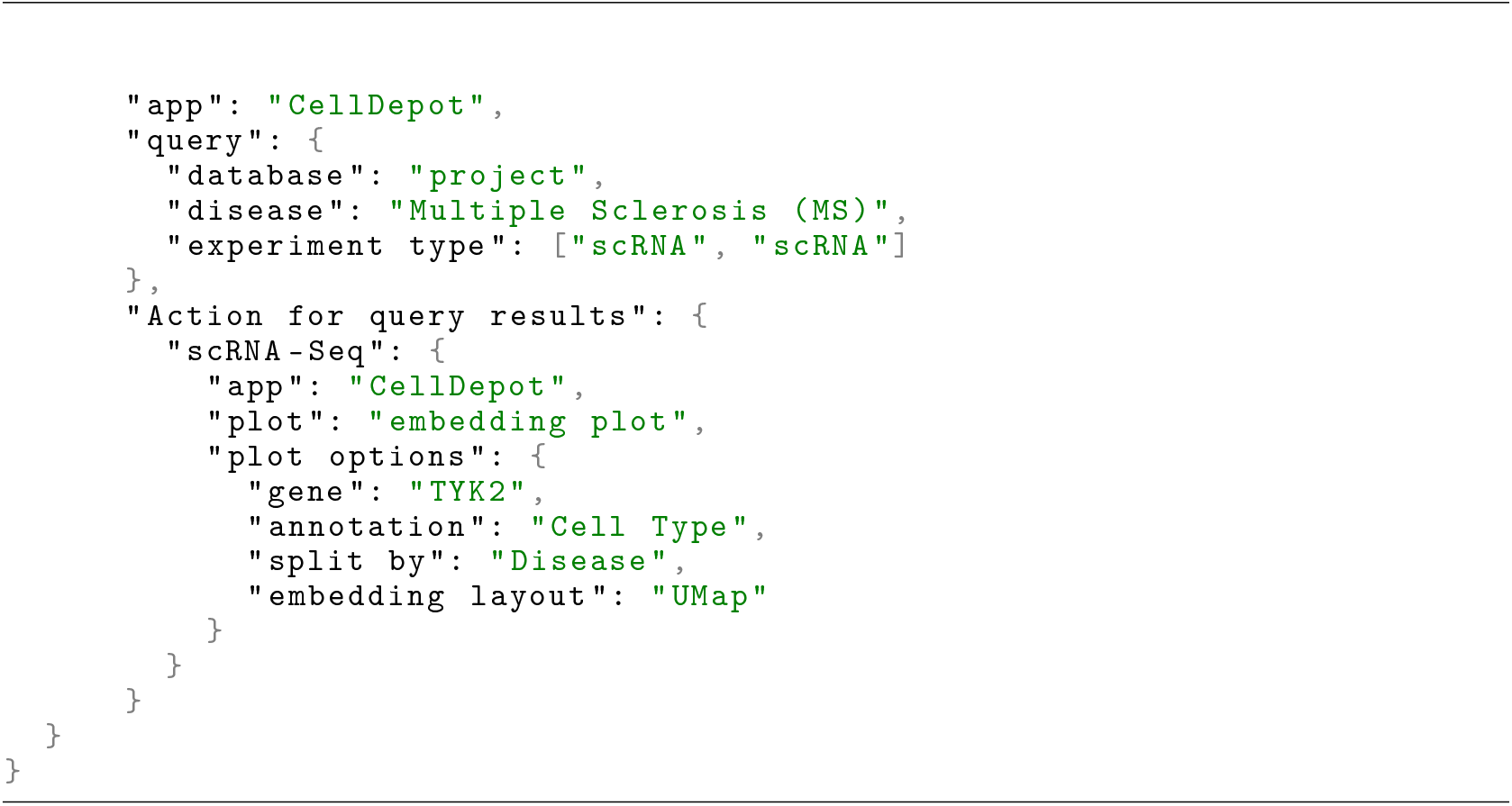

This example illustrates a query to explore the Multiple Sclerosis (MS) dataset and visualize TYK2 gene expression using a UMAP embedding plot. The plot is intended to display the gene’s expression patterns across different cell types, using the annotation of cell types present in the dataset. The data is split by disease conditions, allowing a comparison of TYK2 expression between MS patients and control groups.

### 3.2 Visualization

CompBioAgent, in conjunction with CellDepot and Cellxgene VIP, enables seamless exploration and visualization of single-cell RNA-seq data. We evaluated its effectiveness in five application cases, demonstrating its ability to generate diverse visualizations like UMAP/t-SNE embeddings, violin plots, heatmaps, etc., based on user queries.

#### Case 1: Gene Expression Analysis

As shown in Figure 2, CompBioAgent allows users to query the expression of a specific genes, such as TYK2, within a disease context, such as multiple sclerosis. CompBioAgent retrieves the relevant single-cell RNA-seq data from CellDepot, performs necessary filtering, and generates violin plots comparing the expression of the gene in disease versus control states. This enables scientists to understand how a gene of interest behaves in relation to a specific disease, offering insights into potential biomarkers or therapeutic targets.

**Figure 1:**
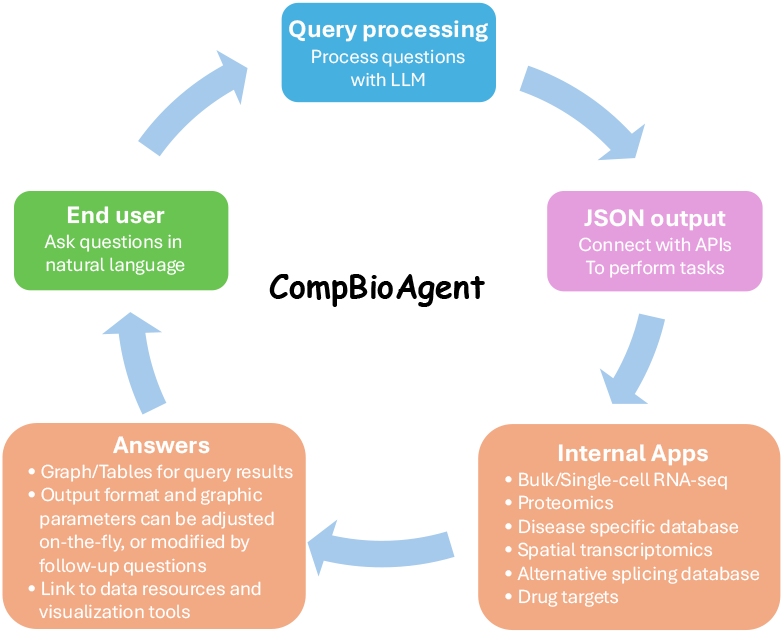
Overview of CompBioAgent.

**Figure 2:**
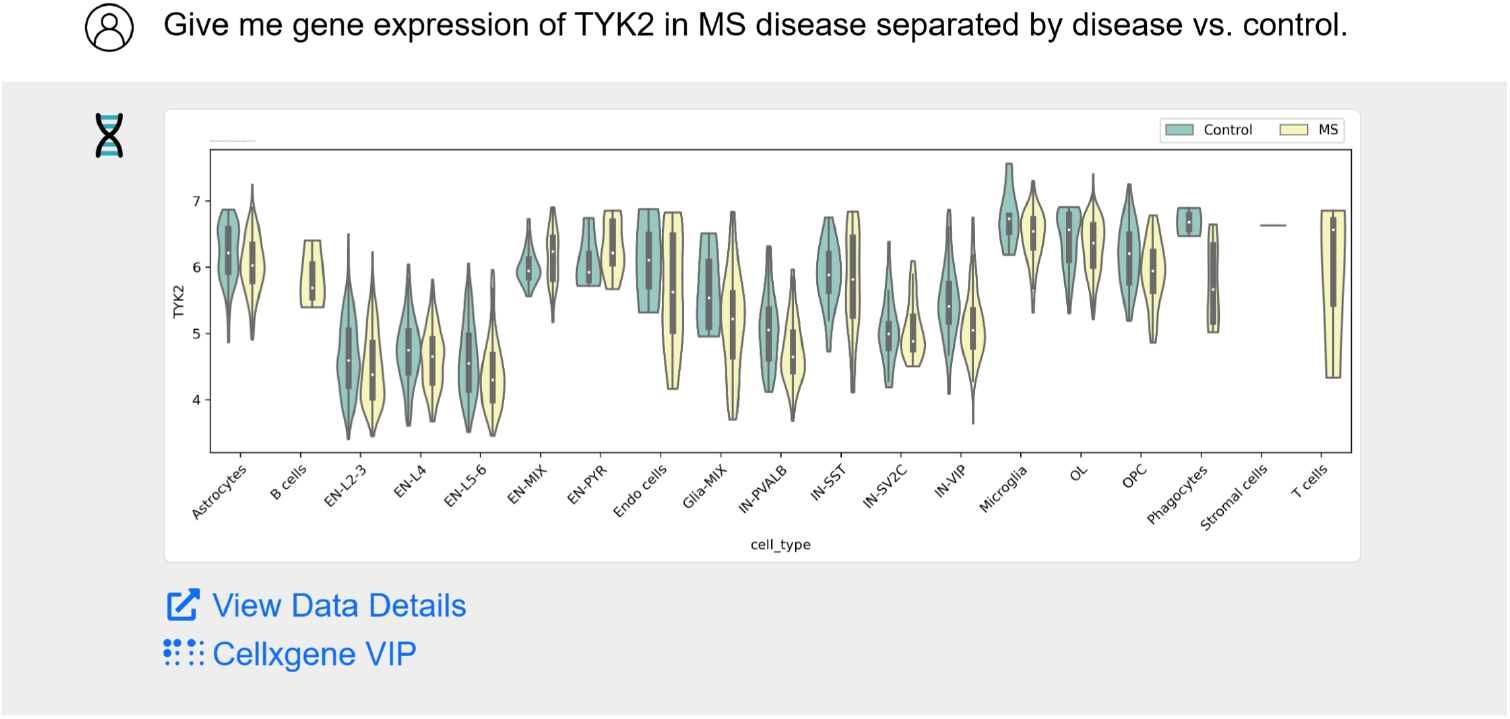
Querying gene expression data for a specific gene (e.g., TYK2) in different disease contexts (e.g., MS), comparing disease versus control conditions (as described in Case 1).

#### Case 2: Heatmap Generation

As shown in Figure 3, CompBioAgent generates heatmaps to display the expression of multiple genes (HLA-B8, SAMHD1, PECAM1, KRT1, CD3D, VIM, MYH11) across CLE samples. CompBioAgent groups cells by disease condition and cell type, creating a visual representation of gene expression patterns across these groups. The heatmap allows researchers to quickly spot patterns or variations in gene expression, facilitating insights into how genes might be differentially regulated in disease contexts. This type of visualization is particularly useful for highlighting gene expression clusters that could be linked to disease progression or specific cellular responses in CLE.

**Figure 3:**
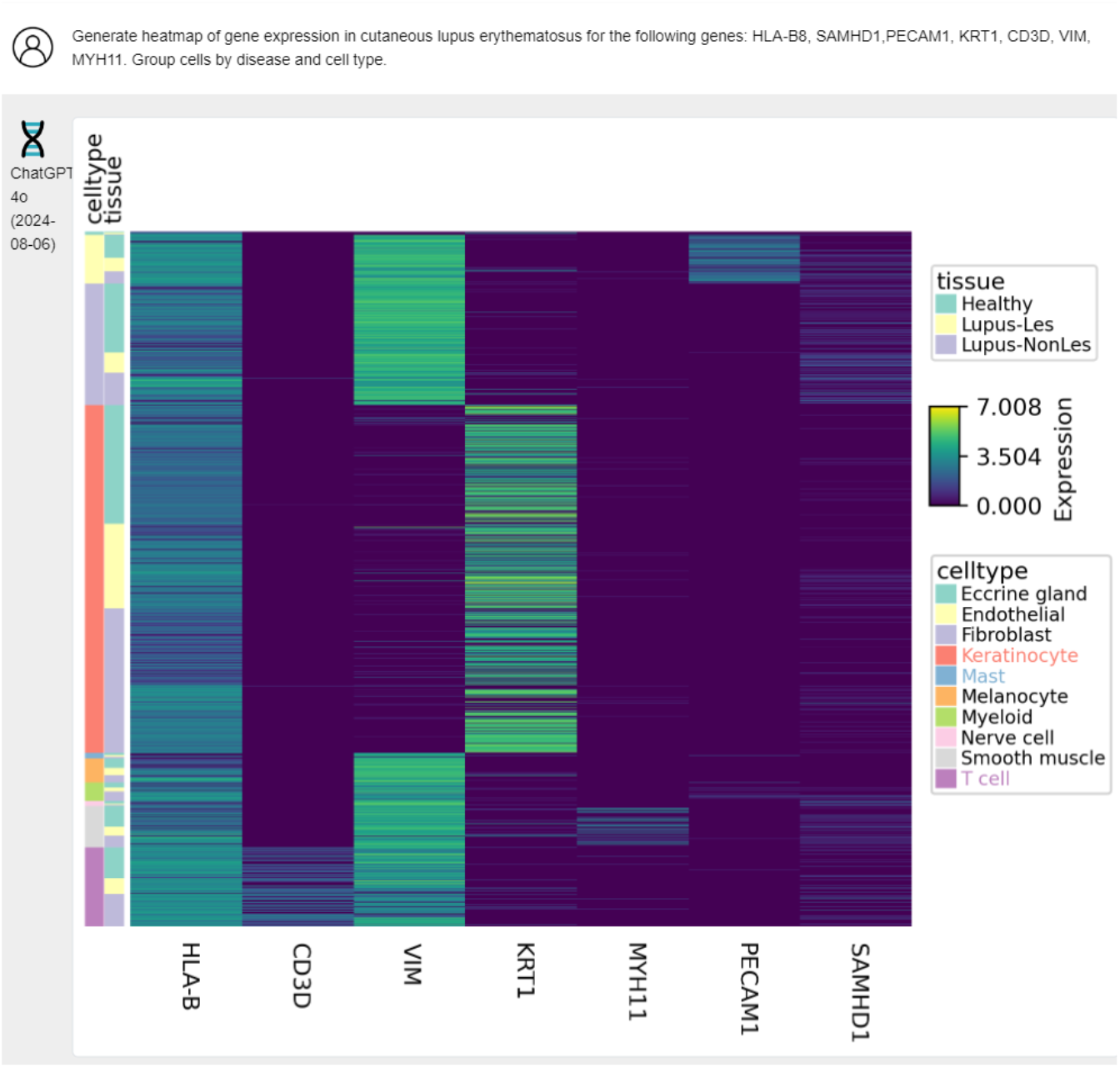
Creating heatmaps for gene expression across multiple genes in CLE, grouping cells by disease and cell type for better insights into gene activity (as described in Case 2).

#### Case 3: Cell Type Proportions

Here, CompBioAgent calculates and visualizes the relative proportions of various cell types in different disease conditions (PPMS, SPMS, Control), as shown in Figure 4. After querying CellDepot for the relevant data, it automatically identifies cell types and generates bar plots to represent the proportions of each cell type across disease and control states. This is crucial for understanding how disease conditions, like MS, affect cellular composition within tissues and how the cell proportion shifts from healthy control to the disease state.

**Figure 4:**
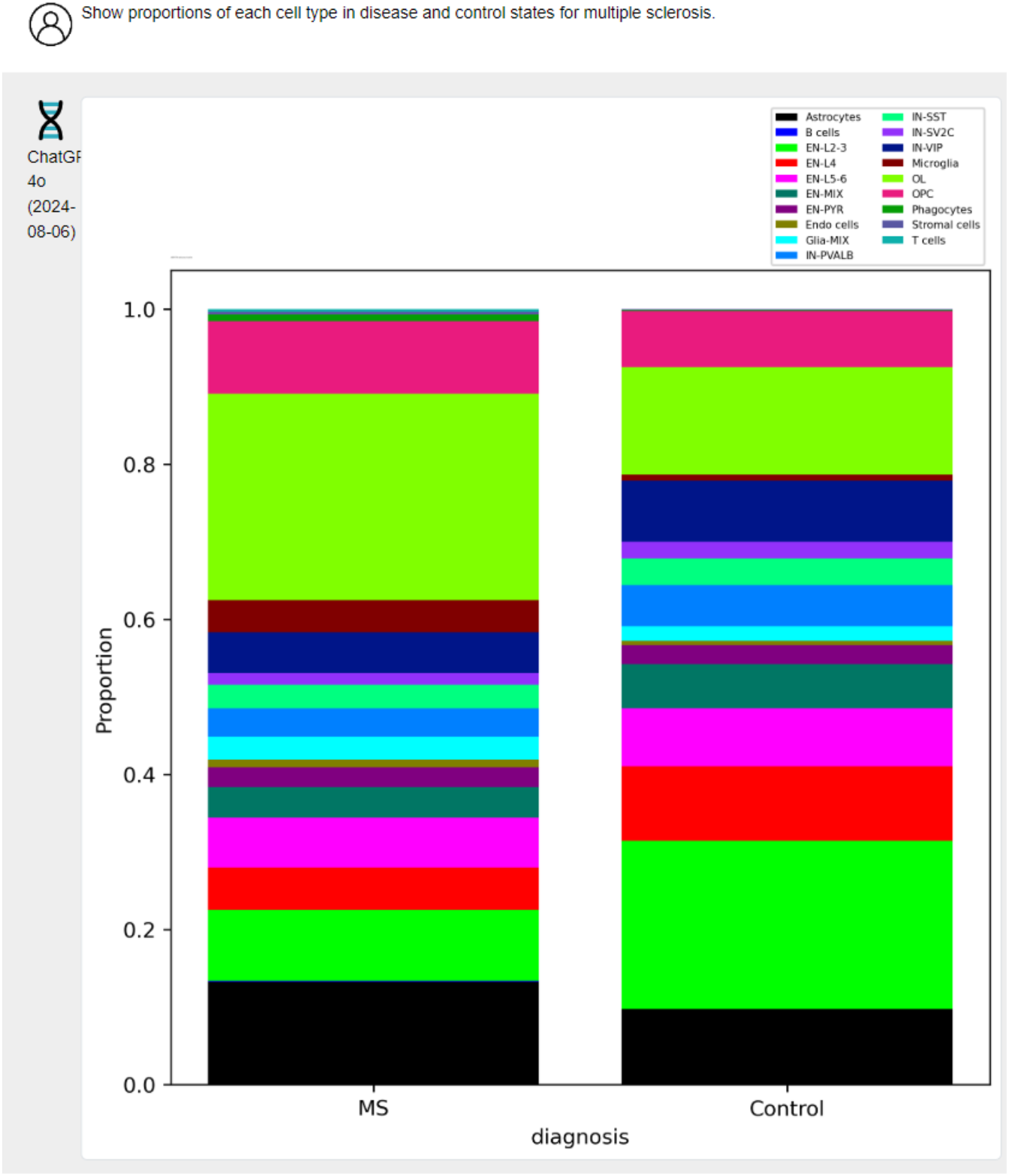
Analyzing the proportion of various cell types in different disease states (e.g., MS disease vs. control) (as described in Case 3).

#### Case 4: Multi-Gene Expression Analysis

This use case involves visualizing the expression of the top 10 marker genes related to cell cycle in different cell types for Glaucoma disease as shown in Figure 5. CompBioAgent queries the relevant datasets from CellDepot and processes the data to exclude any cells with an unknown disease state. Subsequently, it generates dot plots that display gene expression across various cell types, helping researchers identify patterns or co-expressions between genes that might be involved in disease processes. This kind of analysis is essential for understanding the molecular mechanisms underlying diseases and identifying potential therapeutic targets.

**Figure 5:**
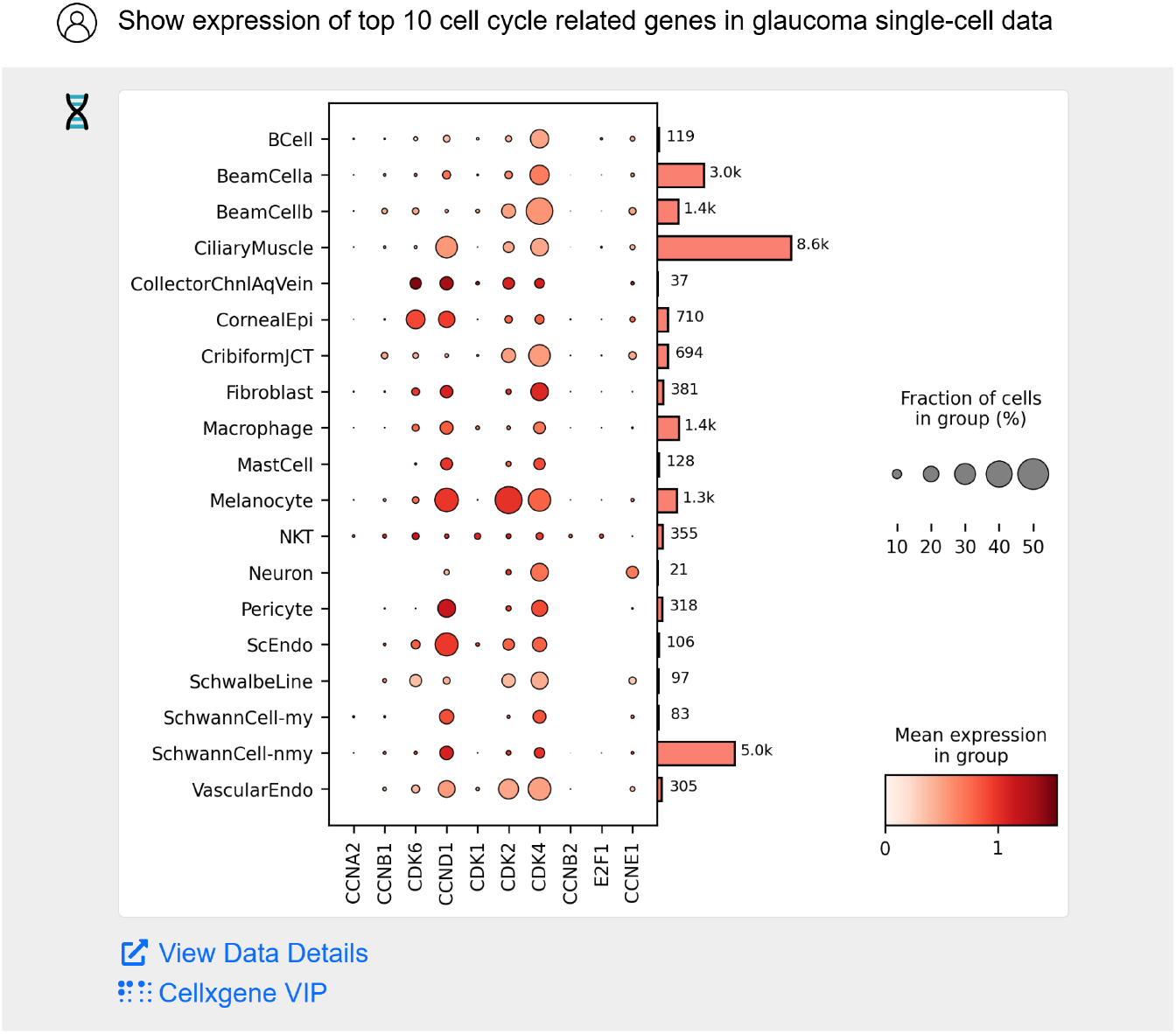
Visualizing gene expression across various cell types for multiple marker genes (e.g., CCND1, CDK2, E2F1, etc.) in diseases like Glaucoma disease, ensuring exclusion of unknown disease states (as described in Case 4).

#### Case 5: UMAP Visualization

As shown in Figure 6, CompBioAgent generates UMAP plots based on the expression of a specific gene, such as TYK2, in multiple sclerosis patients or healthy controls. The plots are automatically split by disease condition (disease vs. control) and are annotated by cell type. CompBioAgent leverages CellxgeneVIP to generate these high-resolution visualizations in real-time. This allows researchers to visualize complex relationships between gene expression, disease condition, and different cell types, making it easier to interpret the heterogeneity of cell populations in diseases.

**Figure 6:**
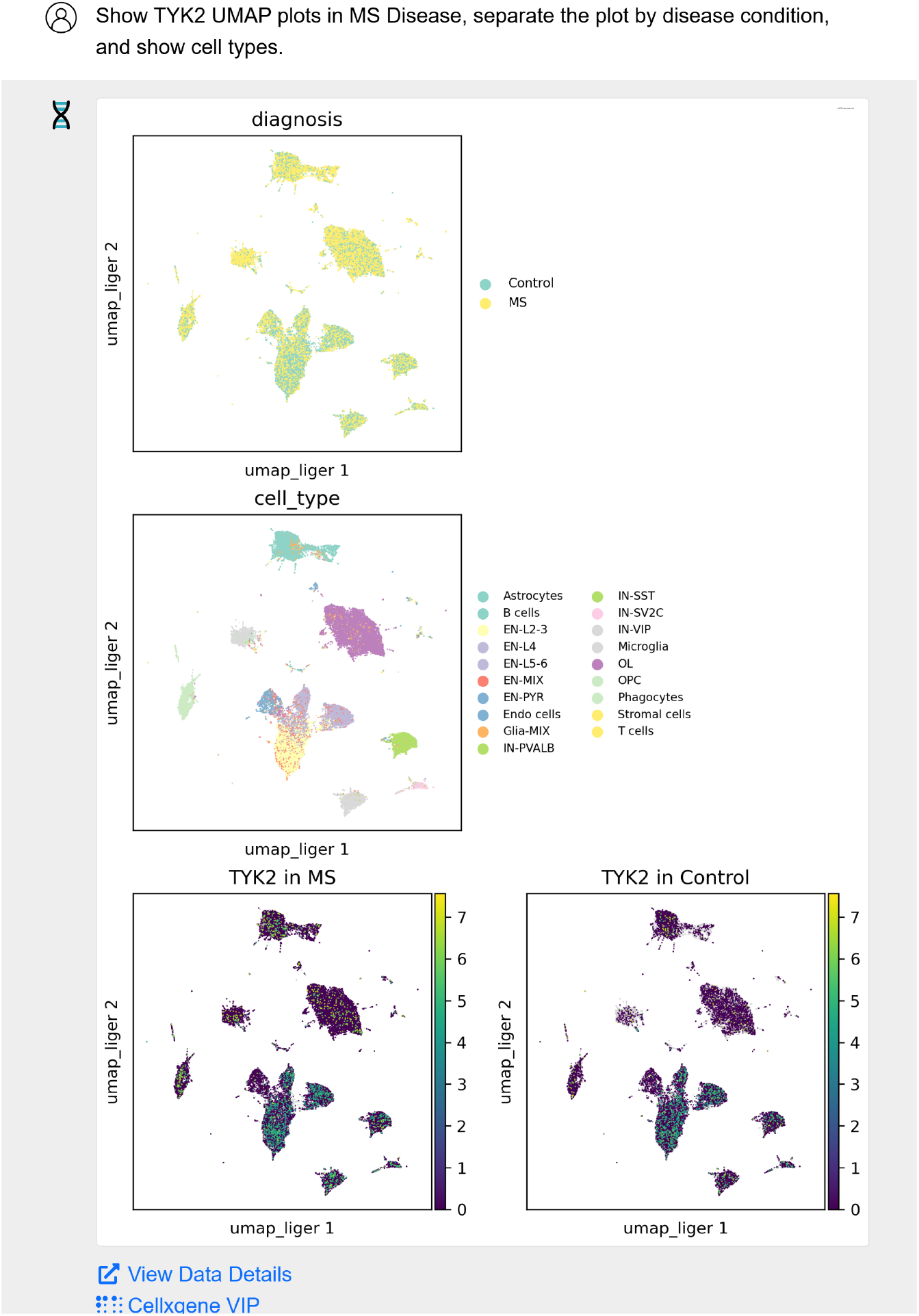
Comparing the expression of key genes in multiple sclerosis (MS) patients and healthy controls across multiple cell types (as described in Case 5).

In all these cases, CompBioAgent acts as a bridge between the user’s natural language input and complex single-cell RNA-seq data analysis. By leveraging the power of Cellxgene VIP and CellDepot, it automatically generates relevant plots, performs necessary data preprocessing (e.g., disease state filtering), and provides interactive, real-time visualizations.

## 4 Conclusion

CompBioAgent represents a significant advancement in making single-cell RNA sequencing (scRNA-seq) data exploration and analysis more accessible and user-friendly. By integrating LLM with platforms like CellDepot and Cellxgene VIP, CompBioAgent allows researchers to efficiently query complex datasets and generate informative visualizations, such as UMAP embedding plots, violin plots, dot plots, heatmaps, etc. This enables users to explore gene expression patterns, analyze cell type distributions, and investigate disease-specific differences, all through natural language that requires no programming expertise.

In this work, we have demonstrated CompBioAgent’s utility in a range of public datasets, particularly in the study of diseases like multiple sclerosis, Alzheimer’s disease, CLE, and others. With its ability to automatically select visualization types and generate plots in real-time, CompBioAgent empowers researchers to quickly derive meaningful insights from scRNA-seq data.

However, there is still room for further development to expand the capabilities of CompBioAgent. Future work may include integrating additional databases and datasets, which will provide researchers with wider range of resources for their analyses. Additionally, incorporating more complex visualizations and plotting options will further enhance the capability of this tool. One important area for improvement is the integration of text-based feedback based on generated plots. This would help users interpret their visualizations by providing context and explanations, enabling more insightful analyses and fostering a deeper understanding of the data.

## References

[1] Fuchoux Tang, Catalin Barbacioru, Yangzhou Wang, Ellen Nordman, Clarence Lee, Nanlan Xu, Xiaohui Wang, John Bodeau, Brian B Tuch, Asim Siddiqui, et al. mrna-seq whole-transcriptome analysis of a single cell. Nature methods, 6(5):377–382, 2009.

[2] Yonatan Katzenelenbogen, Fadi Sheban, Adam Yalin, Ido Yofe, Dmitry Svetlichnyy, Diego Adhemar Jaitin, Chamutal Bornstein, Adi Moshe, Hadas Keren-Shaul, Merav Cohen, et al. Coupled scrna-seq and intracellular protein activity reveal an immunosuppressive role of trem2 in cancer. Cell, 182(4):872–885, 2020.

[3] E Madissoon, A Wilbrey-Clark, RJ Miragaia, K Saeb-Parsy, KT Mahbubani, N Georgakopoulos, P Harding, K Polanski, N Huang, K Nowicki-Osuch, et al. scrna-seq assessment of the human lung, spleen, and esophagus tissue stability after cold preservation. Genome biology, 21:1–16, 2020.

[4] Dongdong Lin, Yirui Chen, Soumya Negi, Derrick Cheng, Zhengyu Ouyang, David Sexton, Kejie Li, and Baohong Zhang. Celldepot: A unified repository for scrna-seq data and visual exploration. Journal of molecular biology, 434(11):167425, 2022.

[5] Leyla Tarhan, Jon Bistline, Jean Chang, Bryan Galloway, Emily Hanna, and Eric Weitz. Single cell portal: an interactive home for single-cell genomics data. BioRxiv, 2023.

[6] Aviv Regev, Sarah A Teichmann, Eric S Lander, Ido Amit, Christophe Benoist, Ewan Birney, Bernd Bodenmiller, Peter Campbell, Piero Carninci, Menna Clatworthy, et al. The human cell atlas. elife, 6:e27041, 2017.

[7] Irene Papatheodorou, Pablo Moreno, Jonathan Manning, Alfonso Muñoz-Pomer Fuentes, Nancy George, Silvie Fexova, Nuno A Fonseca, Anja Füllgrabe, Matthew Green, Ni Huang, et al. Expression atlas update: from tissues to single cells. Nucleic acids research, 48(D1):D77–D83, 2020.

[8] F Alexander Wolf, Philipp Angerer, and Fabian J Theis. Scanpy: large-scale single-cell gene expression data analysis. Genome biology, 19:1–5, 2018.

[9] Severin Dicks, Philipp A., Jura Pintar, Till Korten, Amit Kumar, Isaac Virshup, and Patrick Metzger. scverse/rapids_singlecell: v0.10.6, June 2024.

[10] Wenxing Hu, Haotian Zhang, Yu H Sun, Shaolong Cao, Jake Gagnon, Zhengyu Ouyang, Yuka Moroishi, and Baohong Zhang. Scalesc: A superfast and scalable single cell rna-seq data analysis pipeline powered by gpu. bioRxiv, pages 2025–01, 2025.

[11] Colin Megill, Bruce Martin, Charlotte Weaver, Sidney Bell, Lia Prins, Seve Badajoz, Brian McCandless, Angela Oliveira Pisco, Marcus Kinsella, Fiona Griffin, et al. Cellxgene: a performant, scalable exploration platform for high dimensional sparse matrices. BioRxiv, pages 2021–04, 2021.

[12] Kejie Li, Zhengyu Ouyang, Dongdong Lin, Michael Mingueneau, Will Chen, David Sexton, and Baohong Zhang. cellxgene vip unleashes full power of interactive visualization, plotting and analysis of scrna-seq data in the scale of millions of cells. BioRxiv, 10(2020.08):28–270652, 2020.

[13] Ashish Vaswani, Noam Shazeer, Niki Parmar, Jakob Uszkoreit, Llion Jones, Aidan N Gomez, Łukasz Kaiser, and Illia Polosukhin. Attention is all you need. Advances in neural information processing systems, 30, 2017.

[14] Arun James Thirunavukarasu, Darren Shu Jeng Ting, Kabilan Elangovan, Laura Gutierrez, Ting Fang Tan, and Daniel Shu Wei Ting. Large language models in medicine. Nature medicine, 29(8):1930–1940, 2023.

[15] Alec Radford, Karthik Narasimhan, Tim Salimans, Ilya Sutskever, et al. Improving language understanding by generative pre-training. 2018.

[16] Hugo Touvron, Thibaut Lavril, Gautier Izacard, Xavier Martinet, Marie-Anne Lachaux, Timothée Lacroix, Baptiste Rozière, Naman Goyal, Eric Hambro, Faisal Azhar, et al. Llama: Open and efficient foundation language models. arXiv preprint 2302.13971, 2023.

[17] Yen-Chun Lu, Ashley Varghese, Rahul Nahar, Hao Chen, Kunming Shao, Xiaoping Bao, and Can Li. scchat: A large language model-powered co-pilot for contextualized single-cell rna sequencing analysis. bioRxiv, pages 2024–10, 2024.

[18] Houcheng Su, Weicai Long, and Yanlin Zhang. Biomaster: Multi-agent system for automated bioinformatics analysis workflow. bioRxiv, pages 2025–01, 2025.

[19] Sebastian Lobentanzer, Shaohong Feng, Noah Bruderer, Andreas Maier, Cankun Wang, Jan Baumbach, Jorge Abreu-Vicente, Nils Krehl, Qin Ma, et al. A platform for the biomedical application of large language models. Nature Biotechnology, pages 1–4, 2025.

[20] Juexiao Zhou, Bin Zhang, Guowei Li, Xiuying Chen, Haoyang Li, Xiaopeng Xu, Siyuan Chen, Wenjia He, Chencheng Xu, Liwei Liu, et al. An ai agent for fully automated multi-omic analyses. Advanced Science, 11(44):2407094, 2024.

[21] Nikita Mehandru, Amanda K Hall, Olesya Melnichenko, Yulia Dubinina, Daniel Tsirulnikov, David Bamman, Ahmed Alaa, Scott Saponas, and Venkat S Malladi. Bioagents: Democratizing bioinformatics analysis with multi-agent systems. arXiv preprint 2501.06314, 2025.

[22] Tom B. Brown, Benjamin Mann, Nick Ryder, Melanie Subbiah, Jared Kaplan, Prafulla Dhariwal, Arvind Neelakantan, Pranav Shyam, Girish Sastry, Amanda Askell, Sandhini Agarwal, Ariel Herbert-Voss, Gretchen Krueger, Tom Henighan, Rewon Child, Aditya Ramesh, Daniel M. Ziegler, Jeffrey Wu, Clemens Winter, Christopher Hesse, Mark Chen, Eric Sigler, Mateusz Litwin, Scott Gray, Benjamin Chess, Jack Clark, Christopher Berner, Sam McCandlish, Alec Radford, Ilya Sutskever, and Dario Amodei. Language models are few-shot learners. CoRR, abs/2005.14165, 2020.

[23] Laria Reynolds and Kyle McDonell. Prompt programming for large language models: Beyond the few-shot paradigm. CoRR, abs/2102.07350, 2021.

[24] Lucas Schirmer, Dmitry Velmeshev, Staffan Holmqvist, Max Kaufmann, Sebastian Werneburg, Diane Jung, Stephanie Vistnes, John H Stockley, Adam Young, Maike Steindel, et al. Neuronal vulnerability and multilineage diversity in multiple sclerosis. Nature, 573(7772):75–82, 2019.

[25] Tavé van Zyl, Wenjun Yan, Alexi McAdams, Yi-Rong Peng, Karthik Shekhar, Aviv Regev, Dejan Juric, and Joshua R Sanes. Cell atlas of aqueous humor outflow pathways in eyes of humans and four model species provides insight into glaucoma pathogenesis. Proceedings of the National Academy of Sciences, 117(19):10339–10349, 2020.

[26] John A Tadross, Lukas Steuernagel, Georgina KC Dowsett, Katherine A Kentistou, Sofia Lundh, Marta Porniece, Paul Klemm, Kara Rainbow, Henning Hvid, Katarzyna Kania, et al. A comprehensive spatio-cellular map of the human hypothalamus. Nature, pages 1–9, 2025.

[27] Richard K Perez, M Grace Gordon, Meena Subramaniam, Min Cheol Kim, George C Hartoularos, Sasha Targ, Yang Sun, Anton Ogorodnikov, Raymund Bueno, Andrew Lu, et al. Single-cell rna-seq reveals cell type–specific molecular and genetic associations to lupus. Science, 376(6589):eabf1970, 2022.

[28] Mariano I Gabitto, Kyle J Travaglini, Victoria M Rachleff, Eitan S Kaplan, Brian Long, Jeanelle Ariza, Yi Ding, Joseph T Mahoney, Nick Dee, Jeff Goldy, et al. Integrated multimodal cell atlas of alzheimer’s disease. Nature Neuroscience, 27(12):2366–2383, 2024.

[29] Garrett S Dunlap, Allison C Billi, Xianying Xing, Feiyang Ma, Mitra P Maz, Lam C Tsoi, Rachael Wasikowski, Jeffrey B Hodgin, Johann E Gudjonsson, J Michelle Kahlenberg, et al. Single-cell transcriptomics reveals distinct effector profiles of infiltrating t cells in lupus skin and kidney. JCI insight, 7(8):e156341, 2022.

