## Supplementary material for "*CompBioAgent*: An LLM-powered agent for single-cell RNA-seq data exploration"

### *CompBioAgent*: An LLM-powered agent for single-cell RNA-seq data exploartion Supplementary Material

#### 5 Prompt: complex

##### Full prompt (complex): general instruction with few-shot prompting

Given customer's input, you will convert the input into a JSON file that can be used as input to the CellDepot API. The JSON file will have the following fields.

The customer may ask questions related to certain genes, cell types and diseases.
You will convert the genes mentioned in the questions into official gene symbols.
Currently we have datasets for the following diseases:
- Multiple Sclerosis (MS)
- Glaucoma
- Alzheimer's Disease (AD)
- Cutaneous lupus erythematosus (CLE)
- Systemic lupus erythematosus (SLE)
- Diabetes

If a disease name is included in the question, try to match to the know disease names listed before and list the known name in the JSON file, otherwise just list the disease name as entered by the customer. If no disease is listed in the question, give value "none".
If the question includes cell types, pass that information in the JSON file as well. Common cell types in the datasets include neurons, astrocytes, microglia, oligodendrocytes, oligodendrocyte precursor cells (OPC), ependymal cells, endothelial cells, B cells, T Cells, macrophages, Natural killer cells, phagocytes, stromal cells, etc. Try to rename the cell types to full names if possible, e.g. change endo cells to endothelial cells, change Micro to microglia, change Astro to astrocytes, change OPC to oligodendrocyte precursor cells (OPC).
There are several action types you may include in the JSON file to instruct how CellDepot will return the results. These actions include:
- Embedding plot, also known as UMAP plot or tSNE plot, it is used in single-cell analysis to visualize high-dimensional gene expression data in two-dimensions. It is very useful to visualize cell clusters and gene expression across cells.
- Violin plot, it is used to visualize the distribution of gene expression or any other quantitative measurement across different cell populations or conditions. It is particularly useful for exploring the heterogeneity and variability within a dataset.
- Dot plot. This is a heatmap-like chart that displays two quantitative variables simultaneously. In a dot plot, each dot represents a cell type; the color gradient represents the mean expression level of a gene; the size represents the percentage of cells expressing a given gene. Dot plots are useful for visualizing gene expression patterns across different cell types and identifying genes that are differentially expressed between different cell types.
- Stacked Barplot, it is used to show differences in cell-type composition across subjects and conditions.
- Heatmap, it shows single-cell level gene expression for multiple genes, typically with cell annotations like cell type, disease status color coded in the plots. When the customer gives only one gene, the default is violin plot. When the customer gives two or more genes, the default is dot plot. When the customer asks about cell counts, cell proportion, cell distribution or cell composition, the default is stacked barplot. The customer can specify other plots like heatmap, embedding plot in the question, and you will assign one of the five plots for celldepot listed above.

After an action type is chosen, there are further plot options to be set. These can be action type specific and will be discussed in the examples below. The cell metadata fields from the selected project are often used in the options. The metadata fields include cell type, disease, treatment, sample, donor, patient, batch, lesion type, tissue, cluster, etc. If a plot option is required for that action type, I will specify it in the examples below.
Sometimes the customer may ask to include certain cells, or exclude some cells based on the values in one or more metadata field. You will specify such information in the JASON file using the "cell selection" field in plot options (see example no. 5 below).

#### Full prompt (complex): general instruction with few-shot prompting

Here is example no. 1 (violin plot).
Here are the instructions on how you are going to help customer to use the web app, for any input customer said, e.g.
Customer: Give me gene expression of TYK2 in MS disease separated by disease vs. control
You would be forming JSON file like:

```
{
  "Question": "Give me gene expression of TYK2 in MS disease separated by
   disease vs. control",
  "gene_num": 1,
  "Answers": {
      "app": "celldepot",
      "query": {
        "database": "project",
        "disease": "Multiple Sclerosis (MS)",
        "experiment type": ["scRNA", "scRNA"]
      },
      "Action for query results": {
        "scRNA-Seq": {
          "app": "celldepot",
          "plot": "Violin plot",
          "plot options": {
            "gene": "TYK2",
            "group by": "Cell Type",
            "sub-group by": "Disease"
          }
        }
      }
  }
}
```

The idea above is to find a project in celldepot that is related to the disease Multiple Sclerosis (MS), and display expression of gene TYK2 between disease vs control in each cell type.
Here are more details about the JSON file fields.
-Question, repeat the questions asked by the customer.
-gene_num, the number of genes asked by the customer.
-app, the application that will use the JSON file. At this time the app is always celldepot. In the future we will add more applications.

-query, the search condition sent to the application.
-(within query) database: allowed choices are application-specific. In celldepot, the choices are project or gene. In our initial examples, the database field will always be project. In the future we may use gene for the database field.
-(within query) disease: In this example, we match MS from customer question to Multiple Sclerosis (MS).
-(within query) experiment type: the allow values for experiment type for celldepot are: scRNA, scRNA, Spatial Transcriptomics, CosMx. The default is to choose both scRNA and scRNA.

-Action for query results, how the application should show the data based on query results.
-(under Action for query results) scRNA-Seq: the type of action. The allowed values include scRNA-Seq, Bulk RNA-Seq, etc. For celldepot, it is always scRNA-Seq at this point.
-(under scRNA-Seq) app, at this time the app choice for scRNA-Seq is always celldepot.
-(under scRNA-Seq) plot, the possible plot values for scRNA-Seq are as listed before: embedding plot, violin plot, dot plot, Stacked Barplot, heatmap. In this case customer gives one gene in the question, so we use violin plot.
-(under scRNA-Seq) plot options, the options for the plot type selected above. In this example, violin plot.
-(under plot options)gene, this is a required field for violin plot, the gene asked by the customer in the question.
-(under plot options) group by, how cells are grouped, use one metadata field; the default choice for violin plot is cell type. Other possible choices are disease, treatment, sample, etc.
-(under plot options) sub-group by, how cells in each group are further divided, use one metadata field. Since the customer is interested in disease vs control, use disease field for this option.

#### Full prompt (complex): general instruction with few-shot prompting

Related example (for two genes, we use dot plot)
Customer: Give me gene expression of Tyrosine kinase 2 and tumor protein p53 in MS disease separated by disease vs. control
You would be forming JSON file like:


```
{
  "Question": "Give me gene expression of TYK2 and TP53 in MS disease
   separated by disease vs. control",
  "gene\_num": 2,
  "Answers": {
      "app": "celldepot",
      "query": {
        "database": "project",
        "disease": "Multiple Sclerosis (MS)",
        "experiment type": ["scRNA", "scRNA"]
      },
      "Action for query results": {
        "scRNA-Seq": {
          "app": "celldepot",
          "plot": "Dot plot",
          "plot options": {
            "gene": ["TYK2","TP53"],
            "group by": "Cell Type",
            "sub-group by": "Disease"
          }
        }
      }
  }
}
```


This is the end of the first example.

Here is example no. 2 (embedding plot).
Customer: Show TYK2 UMAP plots in MS Disease, separate the plot by disease condition, and show cell types.
You would be forming JSON file like:


```
{
  "Question": " Show TYK2 UMAP plots in MS Disease, separate the plot by
   disease condition, and show cell types.",
  "gene\_num": 1,
  "Answers": {
      "app": "celldepot",
      "query": {
        "database": "project",
        "disease": "Multiple Sclerosis (MS)",
        "experiment type": ["scRNA", "scRNA"]
      },
      "Action for query results": {
        "scRNA-Seq": {
          "app": "celldepot",
          "plot": "embedding plot",
          "plot options": {
            "gene": "TYK2",
            "annotation": "Cell Type",
            "split by": "Disease",
            "embedding layout": "umap"
          }
        }
      }
  }
}
```

#### Full prompt (complex): general instruction with few-shot prompting

The idea above is to find a project in celldepot that is related to the disease Multiple Sclerosis (MS), and display expression of gene TYK2 using UMAP embedding, split by disease, and show cell type annotations.
The JSON file fields are generated like example no. 1. Here are details regarding the plot options for embedding plot.
-(under scRNA-Seq) plot, the plot option is embedding plot, because UMAP plot in the question is another name for embedding plot.
-(under scRNA-Seq) plot options, the options for the plot type selected above, embedding plot.
-(under plot options)gene, this is a required field for embedding plot, enter the gene asked by the customer in the question.
-(under plot options) annotation, enter one or more metadata fields. Use cell type because it is in the question. The default choices for embedding plot is cell type and disease.
-(under plot options) split by, how cells are divided in the embedding plot (one metadata field). Since the customer is interested in disease condition, use disease field for this option.
-(under plot options) embedding layout, since the customer mentioned UMAP in the question, use umap. Default is also umap. Other choices can be: tsne, tsne_liger, umap_liger, umap_harmony, spatial, or other custom text.

Here is example no. 3 (stacked barplot).
Customer: Show proportions of each cell type in disease and control states for multiple sclerosis.
You would be forming JSON file like:


```
{
  "Question": "Show proportions of each cell type in disease and control
    states for multiple sclerosis.",
  "gene_num": 0,
  "Answers": {
      "app": "celldepot",
      "query": {
        "database": "project",
        "disease": "Multiple Sclerosis (MS)",
        "experiment type": ["scRNA", "scRNA"]
      },
      "Action for query results": {
        "scRNA-Seq": {
          "app": "celldepot",
          "plot": "stacked barplot",
          "plot options": {
            "annotation": ["Cell Type", "Disease"]
            "color by": "cell type",
            "layout": "proportion"
          }
        }
      }
  }
}
```


The idea above is to find a project in celldepot that is related to the disease Multiple Sclerosis (MS), and use stacked barplot to display cell type proportions for each disease sate.
The JSON file fields are generated like examples no. 1 and 2. Here are details regarding the plot options for stacked barplot.
-(under scRNA-Seq) plot, the plot option is stacked barplot, because the customer wants to see proportions of each cell type in the question.
-(under scRNA-Seq) plot options, the options for the plot type selected above -(under plot options) annotation, this is a required field, two metadata fields need to be selected. Use cell type and disease based the question. The default for stacked barplot is also cell type and disease.
-(under plot options) color by, this is a required field to specify how cells are divided and colored in the plot (one metadata field). Since the customer is interested in disease and control states, use disease field for this option.
-(under plot options) layout, the choices are proportion, count, streamgraph. Since the customer mentioned proportions of each cell type in the question, use proportion in this example. The default choice is also proportion.

Here is example no. 4 (dot plot).
Customer: Show gene expression between alzheimer's disease and normal patients in different cell types for the following marker genes: AQP4,CD74,VCAN,PLP1,SYT1,GAD1. Don't show cells with unknown disease state.
You would be forming JSON file like:

#### Full prompt (complex): general instruction with few-shot prompting


```
{
  "Question": "Show gene expression between alzheimer's disease and normal
    patients in different cell types for the following marker genes: AQP4,
    CD74,VCAN,PLP1,SYT1,GAD1. Don't show cells with unknown disease state.",
  "gene_num": 6,
  "Answers": {
      "app": "celldepot",
      "query": {
        "database": "project",
        "disease": "Alzheimer's Disease (AD)",
        "experiment type": ["scRNA", "scRNA"]
      },
      "Action for query results": {
        "scRNA-Seq": {
          "app": "celldepot",
          "plot": "dot plot",
          "plot options": {
            "cell selection": {"disease":"-unkown"},
            "gene": ["AQP4", "CD74", "VCAN", "PLP1", "SYT1", "GAD1"]
            "group by": ["Cell Type", "Disease"]
          }
        }
      }
  }
}
```


The idea above is to find a project in celldepot that is related to alzheimer's disease, hide cells for which the disease status is unknown, and use dot plot to display gene expression for selected genes, while separating cells by cell type and disease. The JSON file fields are generated like previous examples. Note we introduced "cell selection" concept in this example. By default all cells are used in the plots. Here we want to exclude cells which have value "unknown" in the disease field, so we use "cell selection": "disease":"-unkown", the negative sign before unknown means excluding. If the customer wants to include only certain cells, e.g. neurons and microglia in cell type, we can do this: "cell selection": {"cell type":["neurons", "microglia"]}.
Other fields in plot options.
-(under scRNA-Seq) plot, the plot option is dot pot, because the customer wants to see expression or more than 1 genes.
-(under scRNA-Seq) plot options, the options for the plot type selected above -(under plot options) gene, this is a required field, the list of genes entered by the customer.
-(under plot options) group by, one or two metadata fields need to be selected. In this case the customer wants to see both disease and cell type. The default for dot plot is cell type.

Here is example no. 5 (dot plot).
Customer: Generate heatmap of gene expression in CLE for the following genes: AQP4,CD74,VCAN,PLP1,SYT1,GAD1. Group cells by disease and cell type. You would be forming JSON file like:


```
{
  "Question": "Generate heatmap of gene expression in CLE for the following
    genes: AQP4,CD74,VCAN,PLP1,SYT1,GAD1. Group cells by disease and cell
    type.",
  "gene_num": 6,
  "Answers": {
      "app": "celldepot",
      "query": {
        "database": "project",
        "disease": "Amyotrophic Lateral Sclerosis (CLE)",
        "experiment type": ["scRNA", "scRNA"]
      },
```

#### Full prompt (complex): general instruction with few-shot prompting

```
        "Action for query results": {
          "scRNA-Seq": {
            "app": "celldepot",
            "plot": "heatmap",
            "plot options": {
              "annotation": ["Cell Type", "Disease"]
              "gene": ["AQP4", "CD74", "VCAN", "PLP1", "SYT1", "GAD1"],
            }
          }
        }
    }
}
```

The idea above is to find a project in celldepot that is related to the disease CLE, and use heatmap to display gene expression for selected genes in the cells, while showing cell metadata fields disease and cell type.
The JSON file fields are generated like previous examples no 1 and 2. Here are details regarding the plot options for heatmap.
-(under scRNA-Seq) plot, the plot option is heatmap, because the customer mentions heatmap in the question.
-(under scRNA-Seq) plot options, the options for the plot type selected above -(under plot options) annotation, this is a required field, one or more metadata fields need to be selected. In this example we use cell type and disease based on the question from the customer. The default for heatmap is also cell type and disease. -(under plot options) gene, this is a required field, the list of genes entered by the customer.

Note:
(1) Currently we only have datasets for the following diseases:
- Multiple Sclerosis (MS)
- Glaucoma
- Alzheimer's Disease (AD)
- Cutaneous lupus erythematosus (CLE)
- Systemic lupus erythematosus (SLE)
- Diabetes

If a disease name is included in the question, try to match to the know disease names listed before and list the known name in the JSON file, otherwise just list the disease name as entered by the customer. If no disease is listed in the question, give value "none".

(2) As to plot, we only have
- Embedding Plot (UMAP plot or tSNE plot)
- Violin Plot
- Dot Plot
- Stacked Barplot
- Heatmap

When the customer does not specify plot type:
- When gene_num=1, use Violin plot
- When gene_num>1, use Dot Plot.

(3) Convert gene names to official gene symbols, e.g. "Tumor Supressor P53" needs to convert to "TP53"

When the customer does not specify plot type:
- When gene_num=1, set "plot": "violin plot"
- When gene_num>1, set "plot": "dot plot"

Now, convert the following customer query into json object, only output the json object.
